# The YARS Tyrosyl tRNA Synthetase Regulates Senescence Induction and Escape Through the Transcriptional Control of LIN9, a Member of the DREAM Complex

**DOI:** 10.1101/2023.12.19.571263

**Authors:** Hugo Coquelet, Geraldine Leman, Amine Maarouf, Coralie Petit, Bertrand Toutain, Cécile Henry, Alice Boissard, Catherine Guette, Eric Lelièvre, Pierre Vidi, Jordan Guillon, Olivier Coqueret

## Abstract

Senescence is a tumor suppressor mechanism triggered by oncogene expression and chemotherapy treatment. It orchestrates a definitive cessation of cell proliferation through the activation of the p53-p21 and p16-Rb pathways, coupled with the compaction of proliferative genes within heterochromatin regions. Some cancer cells have the ability to elude this proliferative arrest but the signaling pathways involved in circumventing senescence remain to be characterized. We have recently described that malignant cells capable of evading senescence have an increased expression of specific tRNAs, such as tRNA-Leu-CAA and tRNA-Tyr-GTA, alongside the activation of their corresponding tRNA ligases, namely LARS and YARS. We have previously shown that YARS promotes senescence escape by activating proliferation and cell cycle genes but its functions during this proliferative arrest remain largely unknown. In this study, we have continued to characterize the functions of YARS, describing non-canonical transcriptional functions of the ligase. Our results show that YARS is present in the nucleus of growing and senescent cells and interacts with the Trim28 transcriptional regulator. Importantly, YARS binds to the LIN9 promoter, a critical member of the Dream complex responsible for regulating cell cycle transcription. YARS facilitates the binding and the phosphorylation of the type II RNA polymerase and promotes the deposit of activating epigenetic marks on the LIN9 promoter. Consequently, during senescence escape, YARS promotes LIN9 expression and both proteins are necessary to induce the proliferation of emergent cells. These results underscore the unconventional transcriptional functions of YARS in promoting senescence escape by activating LIN9 and likely influencing the functions of the Dream complex.

## INTRODUCTION

Activated in response to various triggers such as oncogenes or chemotherapy (1–3), senescence is a critical biological process that restrains the proliferation of abnormal cancer cells. This tumor suppressor mechanism is generally initiated by sub-lethal DNA damage which then activates the p53-p21 and p16-Rb pathways. A definitive halt in cell proliferation is maintained by the tri-methylation of the lysine 9 of histone H3 which leads to a permanent compaction of proliferative genes (4).

Although senescence induces most of the time a definitive proliferative arrest, we and others have described that this pathway induces an heterogeneous response. For instance, it is well known that the inactivation of the p16INK4 gene disables senescence. Its stability also relies on the dynamic regulation of epigenetic marks. The abnormal up-regulation of the LSD1 and JMJD2C demethylases removes the H3K9 repressive marks and allows cancer cells to escape senescence (5). We have also described that sub-populations of cancer cells can escape chemotherapy-induced senescence (CIS) (3,6–8). Cells capable of evading CIS are more transformed, they resist anoïkis, are more invasive and express a reduced expression of the CD47 receptor (9–12). Recent work using single cell RNA sequencing extended these observations in primary cells, identifying a population of senescence-evading cells in lung fibroblasts treated with doxorubicin (13). Senescence is therefore much more heterogeneous than initially expected and this pathway can function as an adaptive mechanism that allows cancer cells to tolerate chemotherapy treatments (7,8).

Little is known about senescence escape and a better characterization of these arrested cells is necessary since chemotherapy resistance is a major concern. We have recently reported that cells that emerge from senescence express specific tRNAs such as tRNA-Leu-CAA and tRNA-Tyr-GTA and their corresponding aminoacyl-tRNA synthetases (AARS), the Leucyl- and Tyrosyl-tRNA ligases (respectively LARS and YARS) (14). In response to chemotherapy treatment, YARS has unexpected functions, activating cell cycle genes to promote senescence escape. As a consequence, its down-regulation reduced the emergence of persistent cells. Previous studies have already reported unconventional roles of specific tRNA ligases during this proliferative arrest. For instance, the seryl-tRNA ligase interacts with the POT1 member of the telomeric shelterin complex to accelerate replicative senescence (15). These results show that AARSs signaling pathways play important roles in the outcome of senescence but their mechanism of action remains to be characterized.

In this study, we pursued our experiments on the non-conventional functions of YARS. Results show that the ligase is present in the nucleus of growing and senescent cells where it interacts with the Trim28 transcriptional regulator to modulate gene expression. Using unbiased analyses, we identified LIN9, a member of the Dream complex (<u>D</u>P, <u>R</u>B [retinoblastoma], <u>E</u>2F, and <u>M</u>uvB), as a main target of YARS. Dream is a regulatory complex involved in the transcriptional control of cell cycle progression. It is composed of the MuvB core (multi vulva class B) which contains the LIN9, LIN37, LIN52, LIN54 and RBBP4 proteins. In association with E2F4/5 and p130/ p107, the MuvB core functions as a transcriptional repressor in the G0 and G1 phases (16). When the MuvB core is associated with B-MYB and FoxM1, Dream functions as a transcriptional activator to promote the progression of the G2/M phase of the cell cycle. The results presented in this study indicate that YARS promotes the recruitment of the type II RNA polymerase and the deposit of activating epigenetic marks on the LIN9 promoter, leading to the up-regulation of this member of the Dream complex. We also observed that the down-regulation of YARS or LIN9 prevents cell proliferation and induces senescence. In addition, both proteins are necessary for the emergence of proliferative cells since their inactivation reduces senescence escape.

Altogether, these results uncovered unexpected transcriptional functions of YARS during senescence. By inducing LIN9 expression, a member of the Dream complex, the ligase allows cell emergence and regulates the outcome of chemotherapy treatment.

## RESULTS

### The inactivation of the YARS tRNA ligase induces senescence in the MCF7 mammary cell line

As stated above, we have recently described that the down-regulation of YARS inhibits senescence escape (14). To explain this result, we speculated that the inhibition of this ligase leads to cell cycle arrest followed by senescence induction. To test this hypothesis, YARS expression was down-regulated by siRNA in growing cells for 96 hr, using either LS174T (colorectal) or MCF7 (mammary) cancer cells. Following YARS inhibition, no effect was observed on LS174T cells. By contrast, this led to an inhibition of MCF7 cell proliferation (Figure 1A). As a control, we analyzed the effect of other tRNA ligases. The same observation was made when LARS was down-regulated but CARS or WARS inhibition had no effect. This indicates that tRNA ligases have heterogeneous functions on cell proliferation. In MCF7 cells, YARS silencing led to an increased number of cells in the G0/G1 phase of the cell cycle and a decreased number of cells in the S and G2/M phases (Figure 1B). No signs of apoptosis were detected (Figure 1C). Western blot analysis confirmed the efficiency of the different siRNAs (Supplementary Figure 1A).

**Figure 1:**
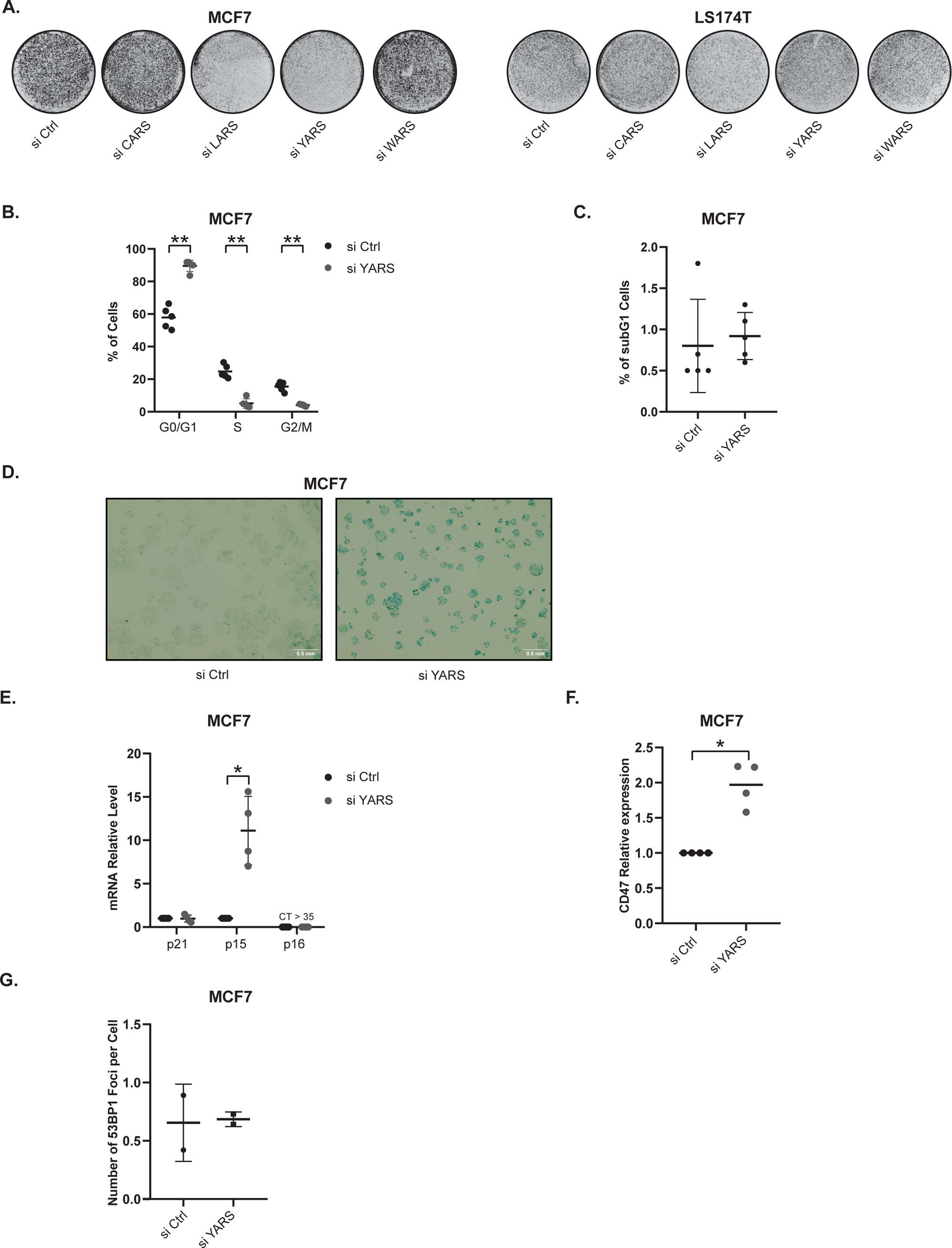
YARS suppression induces senescence in MCF7 cells. A. Growing MCF7 and LS174T cells were transfected with a control siRNA or siRNAs directed against LARS, YARS, WARS or CARS. Cell proliferation was evaluated four days later by crystal violet staining (representative images of experiments for each cell line). B. Growing MCF7 cells were transfected with the indicated siRNAs. After four days, FACS analysis was performed to analyze cell cycle profiles (n=5, ** p<0.01). C. Growing MCF7 cells were transfected with the indicated siRNAs. After four days, FACS analysis was performed to analyze the subG1 population (n=5). D. Representative images of SA-β galactosidase staining, 96 hours after the transfection of MCF7 cells with the indicated siRNAs (n=4). E. Growing MCF7 cells were transfected with the indicated siRNAs. After four days, the expression of p15, p21 and p16 mRNAs was evaluated by RT-qPCR (n=4, * p<0.05). F. Growing MCF7 cells were transfected with a control siRNA or a siRNA directed against YARS. After four days, FACS analysis was performed to analyze CD47 expression (n=4) G. Growing MCF7 cells were transfected with a control siRNA or a siRNA directed against YARS. After fours days, MCF7 cells were fixed and stained with antibody directed against 53BP1. Results were analyzed by fluorescence microscopy and foci counting (n=2).

We then analyzed the main parameters of senescence. YARS inhibition induced a significant increase in ß-galactosidase staining (Figure 1D). This effect was specific since CARS, LARS and WARS had no effect, and this staining was not detected in LS174T colorectal cells (Supplementary Figure 1B). YARS down-regulation did not modify p21 level but significantly increased p15 expression in MCF7 cells but not in LS174T cells (Figure 1E and Supplementary Figure 1C, note that p16INK4 is inactivated in the two cell lines). We and others have recently shown that senescent cells can be identified by an increased expression of the CD47 receptor (12,17). Results presented Figure 1F indicate that YARS inhibition led to a significant increase in the number of CD47^high^ MCF7 cells. As a control, this was also observed when senescence was induced by doxorubicin (Supplementary Figure 1D). Senescent cells are also characterized by an increased DNA damage response. Results presented in Figure 1G indicate that YARS inhibition did not modify the number of 53BP1 foci. As expected, these foci were increased following doxorubicin treatment (Supplementary Figure 1E).

Altogether, these results showed that YARS inhibition led to an induction of senescence in growing MCF7 mammary cells, characterized by an increased ß-galactosidase staining, an up-regulation of p15 expression and the generation of CD47^high^ cells. The effects of tRNA ligases are heterogeneous since YARS inhibition did not induce senescence in LS174T cells and CARS, LARS and WARS inhibition had no effect on senescence induction.

### YARS is present in the nucleus and interacts with the Trim28 transcriptional regulator

tRNA ligase inhibition might lead to the accumulation of translational stress and the consequent induction of cell death. However, no signs of apoptosis were detected in our conditions (see above Figure 1C). In addition, YARS inhibition had no effect on the phosphorylation of eIF2alpha, a common marker activated by uncharged tRNAs and the integrated stress response (Figure 2A, (18,19)). We therefore considered that a significant translational stress was unlikely to explain the induction of senescence in MCF7 cells and searched for a different explanation. Recent results have reported that YARS translocates to the nucleus to regulate gene expression during oxidative stress (20). We therefore determined if the ligase was present in the nucleus in our experimental conditions and if this localization could be related to the regulation of senescence. Using Western blot analysis, we detected YARS in the nuclear extract of growing MCF7 and LS174T cells (Figure 2B). We then determined if this localization could be confirmed by immunofluorescence and if it varies between growing and senescent cells. To this end, LS174T and MCF7 cells were treated respectively with SN38 or doxorubicin, two common drugs used in chemotherapy treatments and well-known inducers of senescence (12,14). CIS induction was confirmed by showing cell cycle arrest, the up-regulation of p21 and p15, SA-β-galactosidase staining, and SASP production (Senescence-Associated Secretory Phenotype, Supplementary Figure 2 A-E). Confocal microscopy experiments and Image J quantification were performed on more than 1200 LS174T nuclei and 2000 MCF7 nuclei. The results presented Figure 2C confirmed the nuclear localization of YARS in growing and senescent cells and indicate that this localization decreased slightly during the proliferative arrest, significantly in MCF7 cells. Control experiments performed in the presence of siRNA directed against YARS resulted in a significant loss of the fluorescence signal (Supplementary Figure 2F).

**Figure 2:**
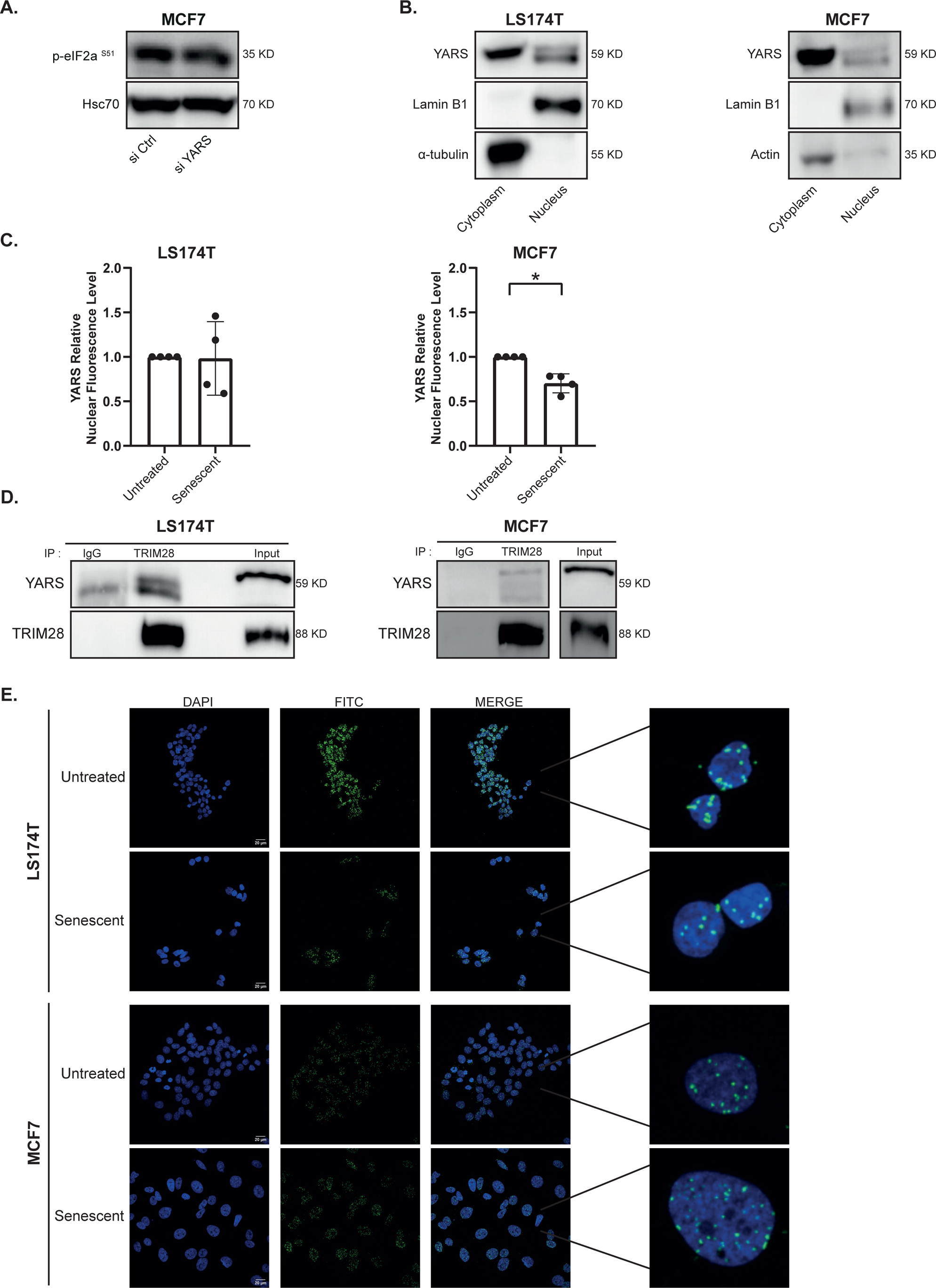
YARS is present in the nucleus and interacts with TRIM28. A. Growing MCF7 cells were transfected with a control siRNA or a siRNA directed against YARS. After three days, eIF2a phosphorylation was analyzed by western blot (n=4). B. Cell extracts were recovered from growing cells, nuclear and cytoplasmic extracts were prepared and the presence of the indicated proteins was analyzed by western blot (n=3 for each cell line). C. Analysis of YARS nuclear expression by confocal immunofluorescence in growing and senescent MCF7 and LS174T cells. YARS nuclear fluorescence intensity was quantified with ImageJ after nucleus segmentation (n=4, * p<0.05). D. Total cell extracts were recovered from growing cells and the association of endogenous TRIM28 and YARS was analyzed by co-immunoprecipitation (n=2 for each cell line). E. TRIM28 and YARS interaction was analyzed by Proximity-Ligation-Assay in growing and senescent cells (representative images of n=4).

These results suggested that YARS might regulate specific nuclear functions in our experimental conditions. During oxidative stress, the ligase interacts with the Trim28 transcriptional regulator to control gene expression (20). To determine if this was the case in our conditions, we first used co-immunoprecipitation experiments and endogenous proteins isolated from MCF7 or LS174T growing cells. Results presented in Figure 2D show that YARS was detected in the two cell lines when Trim28 was immunoprecipitated. When YARS was first immunoprecipitated, Trim28 was again detected (Supplementary Figure 3A). We then used a proximity Ligation Assay (PLA) to confirm this observation in unmodified cells. Confocal microscopy results presented in Figure 2E confirmed the nuclear interaction of YARS and Trim28. This interaction did not vary significantly in senescent cells. Control experiments performed in the presence of siRNAs showed that the down-regulation of YARS resulted in a complete loss of the PLA signal (Supplementary Figure 3B). This signal was also lost when YARS or Trim28 antibodies were replaced by control isotypes (Supplementary Figure 3C).

Altogether, these results indicate that YARS is present in the nucleus of LS174T and MCF7 growing cells and that this nuclear localization is conserved and slightly decreased during chemotherapy-mediated senescence in MCF7 cells. In these two conditions, the ligase interacts with Trim28, suggesting that the ligase might be involved in the regulation of gene transcription.

### LIN9, a member of the Dream complex, is activated by YARS at the mRNA level

We therefore tested the hypothesis that YARS regulates proliferative genes, which would be inhibited in senescent cells and which would contribute to the induction of proliferative arrest. To achieve this, we opted to initially characterize the down-regulated genes common to both cell lines and to the two distinct chemotherapy treatments. Subsequently, we analyzed the potential regulation of selected targets by YARS. To this end, we treated MCF7 and LS174T cells respectively with doxorubicin or sn38 for 96hr and performed proteomics analysis. We identified 1781 common proteins that were down-regulated during senescence induction. GSEA analysis indicated that these proteins corresponded as expected to cell cycle and DNA damage pathways but more significantly to target genes of the Dream complex (pvalue 3.96X10^−198^, Figure 3A). We confirmed these results at the transcriptional level using RNA-seq and ATAQ-Seq experiments in LS174T senescent cells. We focused on the repressed genes whose promoters were more compacted and mRNA expression decreased since senescence pathways are well known to induce the epigenetic compaction of proliferative genes (3,4). This approach identified 3637 genes, among which a significant deregulation of Dream target genes was also detected by GSEA analysis (p-value 3.1X10^−292^, Figure 3B).

**Figure 3:**
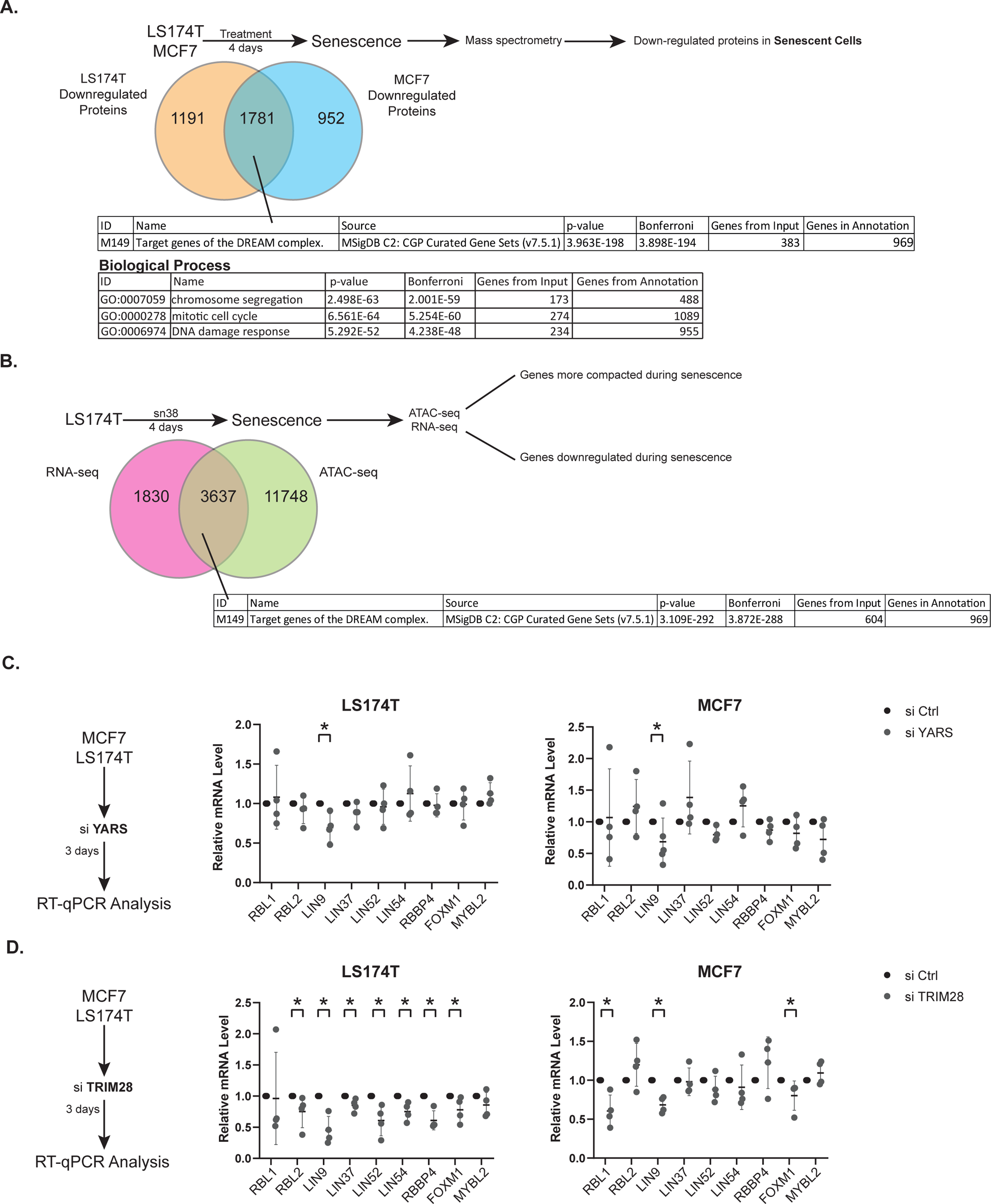
YARS regulates the expression of LIN9, a member of the DREAM complex. A. Senescent LS174T and MCF7 cell extracts were recovered and analyzed by quantitative proteomics (n=3 for each cell line). B. Senescent LS174T cell extracts were recovered and RNA and ATACseq analysis were performed and analyzed with ToppGene (RNAseq n=3, ATAC-seq n=2). C. Growing MCF7 and LS174T cells were transfected with a control siRNA or a siRNA directed against YARS for 24h and cell extracts were recovered 3 days after depletion. The expression of the indicated mRNAs was analyzed by RT-qPCR (n=4 for each cell line, * p<0.05). D. Growing MCF7 and LS174T cells were transfected with a control siRNA, or a siRNA directed against TRIM28 for 24h and cell extracts were recovered 3 days after depletion. The expression of the indicated mRNAs was analyzed by RT-qPCR (n=4 for each cell line, * p<0.05).

These unbiased analyses identified the Dream pathway as a main target inhibited during senescence induction, in two different cell lines and in response to two different treatments. As stated above, Dream is an essential transcriptional regulator of cycle progression (16) and recent results have described that this complex is required for the induction of Ras-mediated senescence (21). We therefore speculated that its expression might be controlled by YARS to regulate the induction of this suppressive arrest. To test this hypothesis, we first investigated if specific members of Dream were regulated by the ligase at the transcriptional level. To this end, YARS was inactivated by siRNA in growing LS174T or MCF7 cells and mRNA expression was analyzed by RT-QPCR after 96hr. Results presented Figure 3C show that the expression of most Dream members was not modified. However, YARS inhibition significantly reduced LIN9 mRNA expression in the two cell lines (Figure 3C). Interestingly, LIN9 level was also reduced when Trim28 was down-regulated by siRNA (Figure 3D, see supplementary Figure 4A for the validation of the Trim28 siRNA).

Taken together, these analyses indicated that the Dream complex or its targets genes are down-regulated in senescent cells. Among its different components, the expression of the LIN9 mRNA was reduced following YARS down-regulation. This suggested that the ligase might control LIN9 expression and Dream functions during senescence. Interestingly, its has already been reported that LIN9 silencing induces senescence in human and mouse fibroblasts (22,23). We therefore focused on this gene and on its regulation by YARS.

### Trim28 and YARS interact with the LIN9 promoter to regulate the binding of the type II RNA polymerase

Since recent studies have reported that YARS can interact with DNA (20), we analyzed its presence on the LIN9 proximal promoter using chromatin immunoprecipitation (ChIP, Figure 4A). Results indicated that YARS was associated with this promoter in dividing cells and that this association was significantly lost in senescent cells (Figure 4B and 4C). These experiments also showed that Trim28 was recruited to this region and that its binding was also reduced during the proliferative arrest. To ensure the specificity of these results, we repeated these experiments following the transfection of control or specific siRNAs targeting Trim28 or YARS. The association of these two proteins with the LIN9 promoter was significantly reduced (Supplementary Figure 4B). We then determined if the binding of YARS was independent of Trim28. To do this, we silenced Trim28 using siRNA and analyzed the interaction of the ligase with the LIN9 promoter. Results presented Figure 4D indicate that Trim28 inhibition decreased YARS DNA binding. Similarly, down-regulating YARS expression led to a reduced interaction of Trim28 with the LIN9 promoter. These results suggested that these two proteins cooperate to interact with this promoter.

**Figure 4:**
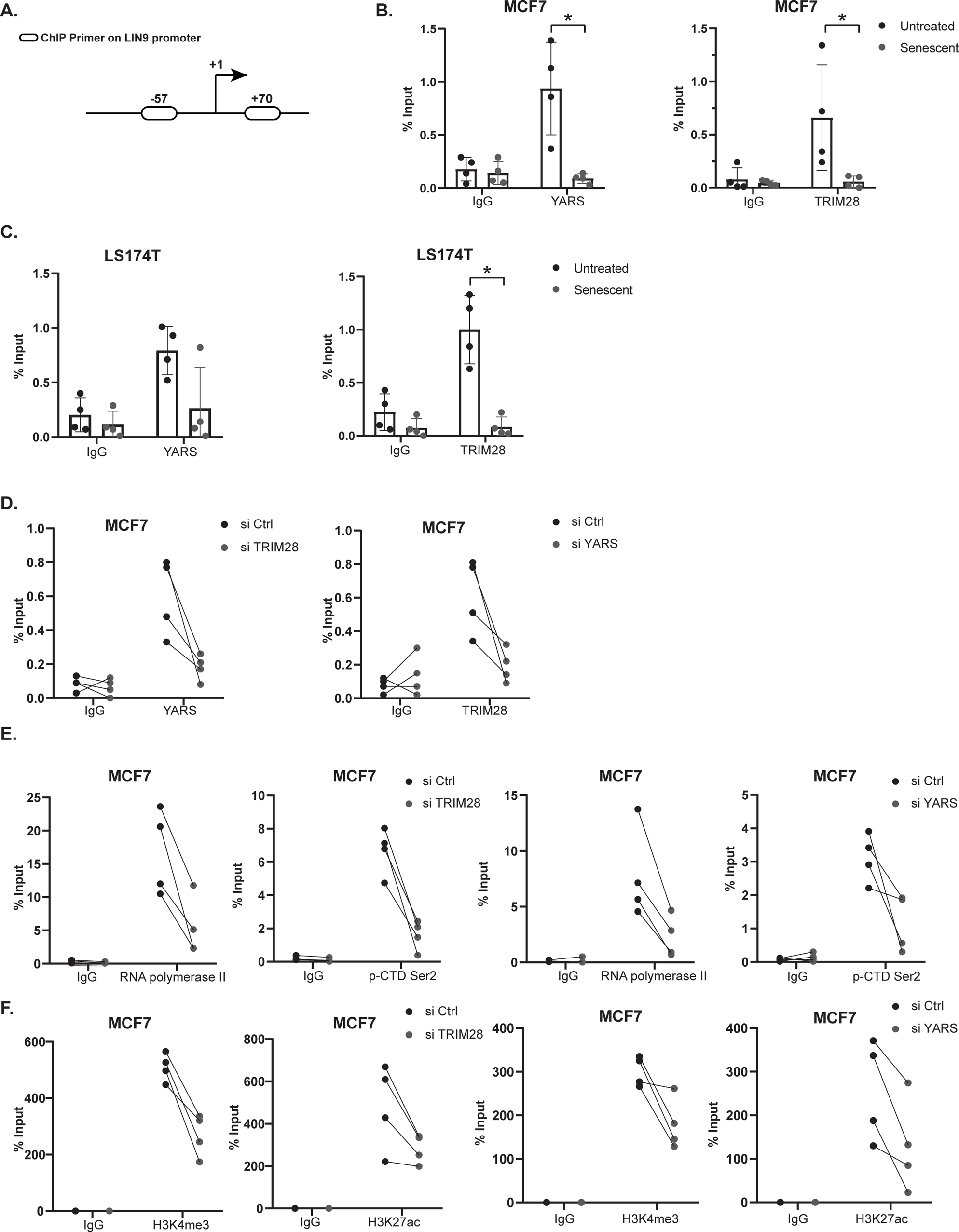
YARS promotes the association of the type II RNA polymerase with the LIN9 promoter to induce its transcriptional activation. A. Schematic view of LIN9 promoter and of the position of the ChIP primers. B and C. Chromatin from growing and senescent MCF7 cells (B) or LS1T4T (C) was recovered and the binding of TRIM28 and YARS to the LIN9 promoter was analyzed by ChIP experiments (n=4 for each cell line, * p<0.05). D. Growing MCF7 cells were transfected with a control siRNA or a siRNA directed against YARS or TRIM28 for 24h. Chromatin was recovered 3 days after depletion, then the binding of the indicated proteins was analyzed by ChIP experiments (n=4, the results for the YARS ChIP following siCtrl are 0.7731, 0.7974, 0.3282, 0.4758 and for the Trim28 ChIP following siCtrl are 0.3409, 0.8141, 0.7836, 0.5101). E. Growing MCF7 cells were transfected with a control siRNA or a siRNA directed against YARS or TRIM28 for 24h. Chromatin was recovered 3 days after depletion, then the binding of the RNA polymerase II and its phosphorylation of the serine 2 residue of the CTD domain were analyzed by ChIP experiments (n=4). F. Growing MCF7 cells were transfected as above in E., chromatin was recovered 3 days after depletion and the presence of the H3K4me3 and H3K27ac epigenetic marks was analyzed by ChIP experiments (n=4).

We then analyzed the transcriptional effects of YARS and Trim28. Trim28 regulates gene elongation by controlling the serine 2 phosphorylation of the C-terminal domain of the type II RNA polymerase (24,25). We confirmed this observation on the LIN9 promoter since the down-regulation of Trim28 by siRNA led to the loss of RNA polymerase binding and consequently of its Ser 2-phosphorylated form (Figure 4E). The same observation was made when YARS expression was inhibited. The reduced expression of the ligase decreased the association of the RNA polymerase and of its phosphorylated form to the LIN9 promoter. We also analyzed the effect of YARS and Trim28 silencing on H3K4 tri-methylation and H3K27 acetylation, two main epigenetic marks of transcriptional activation. Results presented Figure 4F show that the down-regulation of the two proteins reduced the presence of these two marks on the proximal promoter of LIN9.

Altogether, these results indicate that YARS and Trim28 activate LIN9 expression at the transcriptional level. The two proteins are present on the LIN9 promoter to promote the binding of the type II RNA polymerase and the deposit of the H3K4me3 and H3K27ac epigenetic marks. In senescent cells, the association of YARS and Trim28 with the LIN9 promoter is significantly reduced, suggesting that LIN9 expression might be inhibited during the proliferative arrest.

### LIN9 down-regulation induced senescence

In light of the above results, we then analyzed LIN9 expression in senescent cells. RT-QPCR experiments presented Figure 5A show that its level was down-regulated in L174T and MCF7 cells following chemotherapy treatment. To analyze if this inhibition was simply a consequence of the proliferative arrest or if LIN9 was actually involved in senescence induction, we suppressed its expression by siRNA in dividing cells. We confirmed the loss of LIN9 at the mRNA level since we were unable to detect the protein using available antibodies. As an additional control, we confirmed the decreased expression of cell cycle genes involved in the progression of the S and G2 phases, the expected consequence of Dream inactivation (Supplementary Figure 4C and 4D). After 96 hours, LIN9 inactivation resulted in a significant reduction of MCF7 cell proliferation but had no effect in LS174T cells, showing once again the heterogeneity of the responses of these two cell lines. (Figure 5B). In the breast cell line, this was correlated with a slight decrease in the number of cells in the G1 and S phases of the cell cycle and an increased number of cells in the G2/M phase (Figure 5C). No modification of the cell cycle profile was noticed in LS174T cells. Results also showed that LIN9 inactivation led to an increased beta-galactosidase staining in MCF7 cells and the up-regulation of p15 and p21 mRNA levels (Figure 5 D-E).

**Figure 5:**
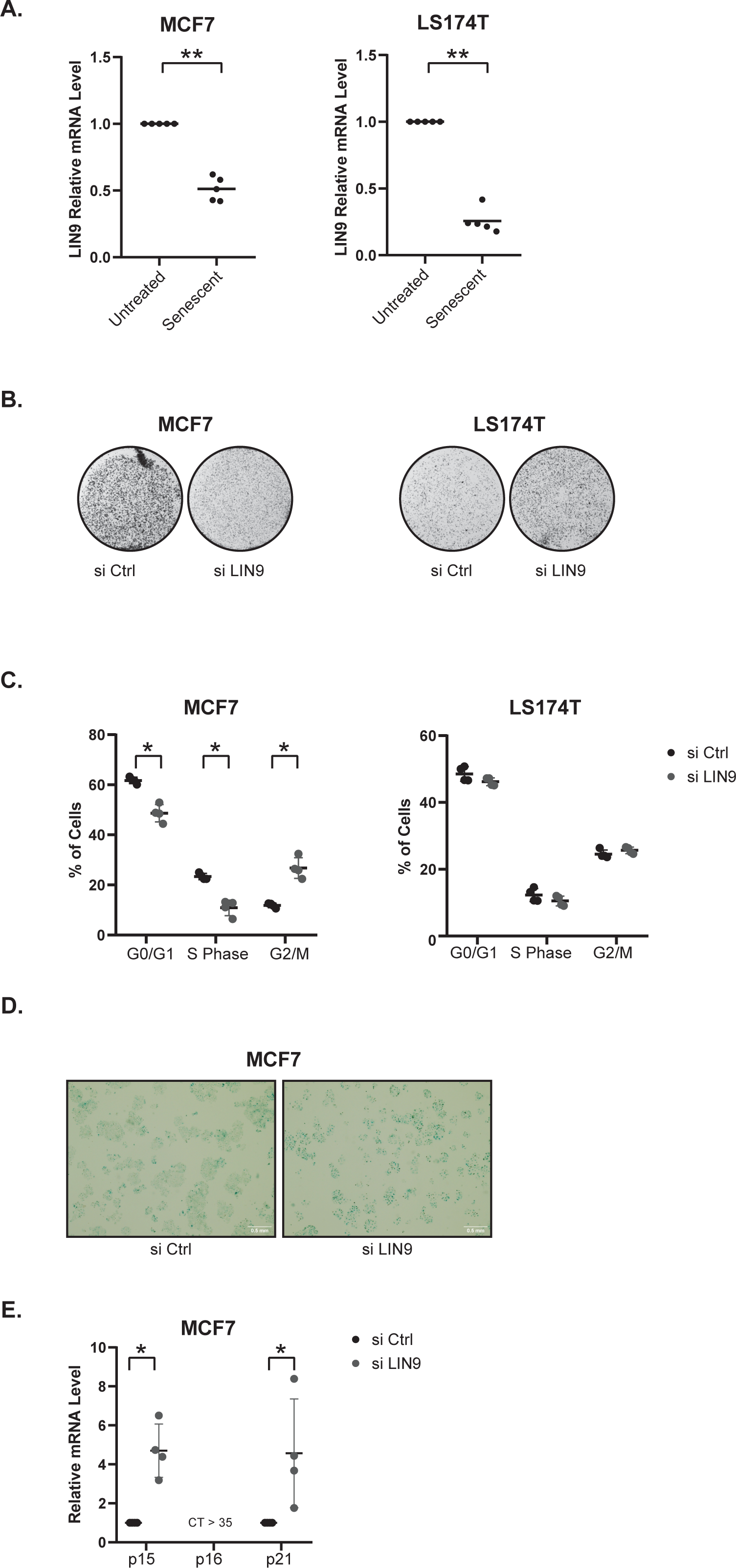
The down-regulation of LIN9 induces senescence in MCF7 cells. A. Growing MCF7 and LS174T cells were treated respectively with doxorubicin or sn38 and cells extracts were recovered after 96hr. The expression of the LIN9 mRNA was analyzed by RT-qPCR (n=5 for each cell line, ** p<0.01). B. Growing MCF7 and LS174T cells were transfected with a control siRNA or a siRNA directed against LIN9 for 24h. Cell proliferation was evaluated 3 days later by crystal violet staining (n=3 for each cell line). C. Growing MCF7 and LS174T cells were transfected as above in B. And after 4 days, FACS analysis was performed to analyze the cell cycle profile of the indicated cells (n=4 for each cell line, * p<0.05). D. Representative images of SA-β galactosidase staining four days after the transfection of MCF7 cells with a control siRNA or an siRNA directed against LIN9 (n=3). E. Growing MCF7 cells were transfected with the indicated siRNAs as described above. After four days, the expression of p15, p21 and p16 mRNA was evaluated by RT-qPCR analysis (n=4, * p<0.05).

Overall, these findings indicate that LIN9 inactivation leads to the induction of senescence in MCF7 cells. This effect was not observed in LS174T cells where YARS inactivation had also no effect on senescence (see above Figure 1). These results suggested that the control of LIN9 expression by YARS might be important during senescence induction.

### The YARS-LIN9 pathway induces CIS escape in MCF7 cells

As stated above, we have previously described that cells can escape chemotherapy-induced senescence and that YARS is involved in the proliferation of emergent cells (14). The above results suggested that the ligase might be present in the nucleus of these emergent cells to control LIN9 expression and promote senescence escape. To test this hypothesis, we first determined if these proteins were involved in senescence escape. Results presented Figure 6A confirmed our recent observation showing that YARS inactivation reduced the emergence of proliferating cells. The same effect was observed when Trim28 was down-regulated (Figure 6B). Note that YARS silencing also reduced senescence escape in LS174T cells even though it did not affect the induction of the proliferative arrest in this cell line (see Supplementary Figure 1B above). This indicates that the ligase controls cell proliferation and emergence by several signaling pathways.

**Figure 6:**
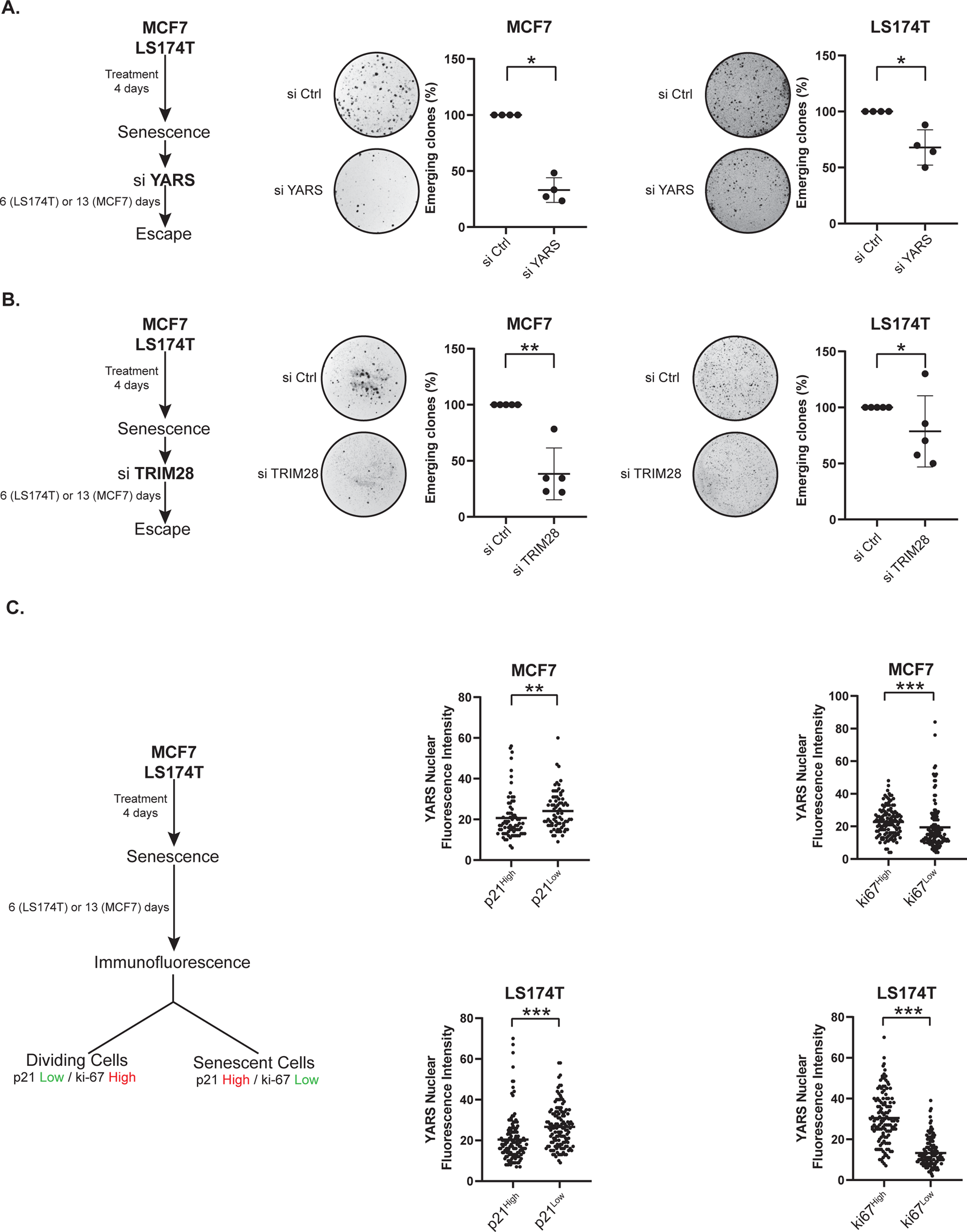
YARS is present in the nucleus of emergent cells and its inactivation reduces senescence escape. A. Senescent MCF7 and LS174T cells were transfected with a control siRNA or a siRNA directed against YARS. The number of emergent cells was evaluated by crystal violet staining 6 days after for LS174T cells or 13 days for MCF7 cells (n=4 for each cell line, * p<0.05). B. Senescent MCF7 and LS174T cells were transfected with a control siRNA or a siRNA directed against TRIM28 for 24h. The number of emergent cells was evaluated by crystal violet staining 6 days after for LS174T cells or 13 days for MCF7 cells (n=5 for each cell line, ** p<0.01, * p<0.05). C. MCF7 and LS174T cells were treated respectively with doxorubicin or sn38 for 96h to induce senescence. Escape was induced by adding 10% fetal bovine serum for 6 days (LS174T) or 13 days (MCF7). Cells were then fixed and stained with antibodies directed against YARS, p21 or ki67 as indicated. Results were analyzed by confocal fluorescence microscopy. The 5% of ki67 or p21 higher and lower expressing cells were selected and the nuclear intensity of YARS staining was analyzed with imageJ (Quantification of around 75 nucleus from 2 different cell culture replicates, ** p<0.01, *** p<0.001).

We then determined if the nuclear expression of the ligase varies during CIS escape. We have recently shown that the dividing cells that escape senescence express the KI67 proliferative antigen and have a reduced expression of p21 (10,12). This population can therefore be identified as KI-67^high^ and p21^low^ cells. Cells that remain senescent are identified as KI-67^low^ and p21^high^ cells (Guillon, Petit et al., 2019). The immunofluorescence experiments presented in Figure 6C indicate that YARS nuclear localization was increased in KI-67^high^ and p21^low^ dividing cells as compared to KI-67^low^ and p21^high^ cells that remain senescent. This observation indicates that the nuclear expression of YARS increases in cells that escape senescence.

We then determine if LIN9 expression was re-activated by YARS in emergent cells. To test this hypothesis we first measured its mRNA level in emergent cells. Results indicate that its expression increased significantly during senescence escape, both in LS174T and MCF7 cells (Figure 7A). When YARS was inactivated by siRNA in senescent cells, a reduced expression of LIN9 mRNA was observed during CIS escape but this was significant only in MCF7 cells (Figure 7B). Next, we explored the potential involvement of LIN9 in senescence escape. As shown in Figure 7C, the inactivation of LIN9 in senescent cells almost completely inhibited CIS escape in MCF7 cells. LIN9 down-regulation only had a slight effect in LS174T cells, further illustrating the difference between the two cell lines.

**Figure 7:**
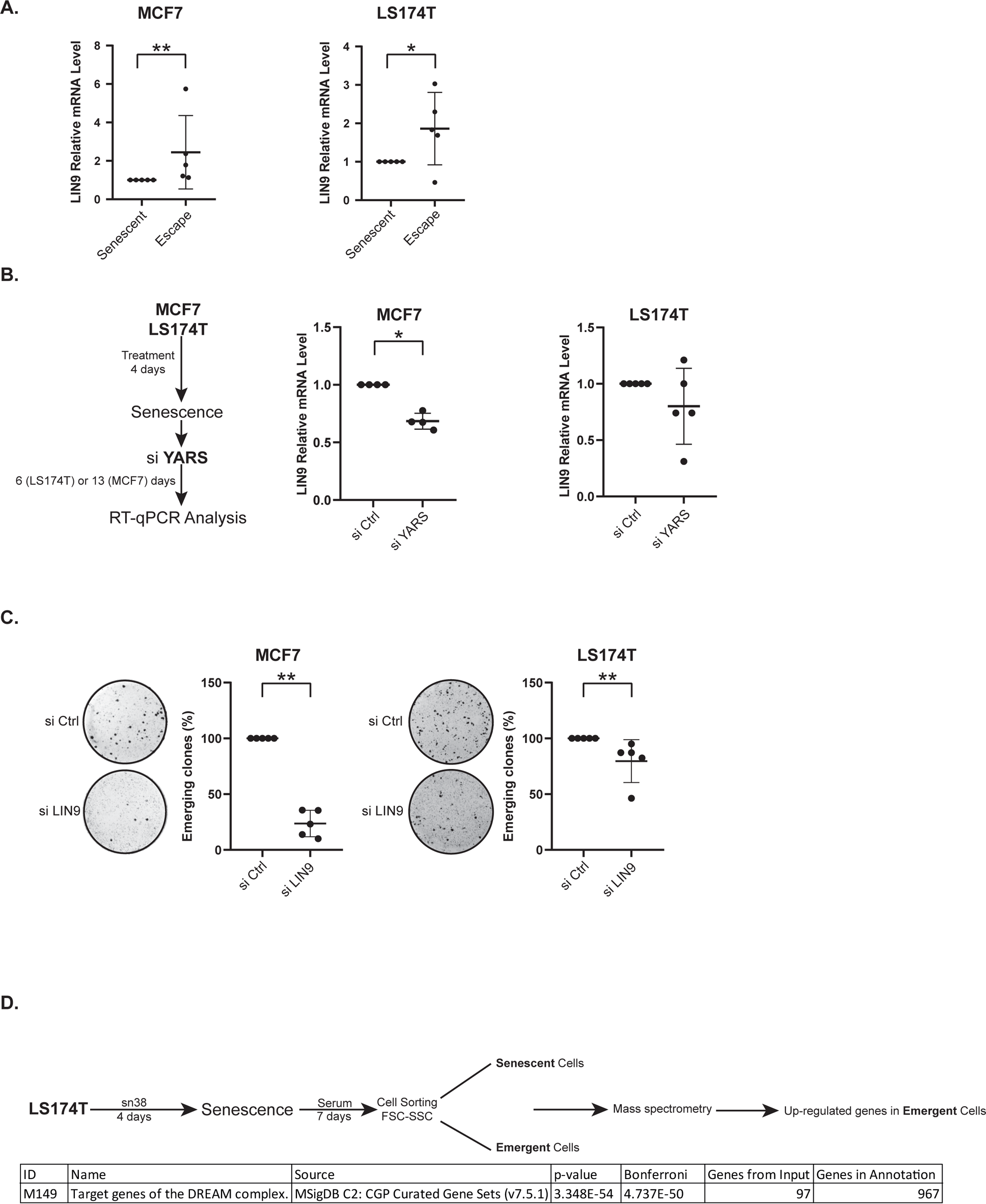
YARS control LIN9 expression during senescence escape. A. Senescent LS174T cells were generated as above and cell emergence was induced by adding 10% fetal bovine serum. After 7 days, cell sorting and enrichment of the dividing (low FSC/SSC) and senescent (high FSC/SSC) populations was performed using flow cytometry as previously described (10,12). Cell extracts were analyzed by SWATH quantitative proteomics and ToppGene analysis (n=3). B. Growing MCF7 and LS174T cells were treated for 96h and escape was induced by by adding 10% fetal bovine serum for 6 days (LS174T) or 13 days (MCF7). Cell extracts were then recovered and the expression of the LIN9 mRNA was analyzed by RT-qPCR (n=5 for each cell line, ** p<0.01 * p<0.05). C. Senescent MCF7 and LS174T cells were transfected with a control siRNA or a siRNA directed against YARS for 24h and escape was induced by adding 10% fetal bovine serum. Cell extracts were recovered 6 days (LS174T) or 13 days (MCF7) after depletion and the expression of the indicated mRNAs was analyzed by RT-qPCR (MCF7 n=4, LS174T n=5, * p<0.05). D. Senescent MCF7 and LS174T cells were transfected with a control siRNA or a siRNA directed against LIN9 for 24h. The number of emergent cells was evaluated 6 or 13 days later (n=5 for each cell line, ** p<0.01).

Since LIN9 expression was reactivated by YARS during cell emergence, we finally determined if a general activation of the Dream complex could be detected during senescence escape. As we previously described (9–11), we used flow cytometry and cell sorting to enrich emerging, dividing cells from the senescent population based on FSC and SSC criteria. Proteomic analysis performed on these two sub-populations showed that the Dream pathway was significantly increased in the emerging, dividing population as compared to cells that remain senescent (p-value 3.34X10-54, Figure 7D).

Altogether, these results indicate that LIN9 expression is necessary for senescence escape and that its expression is re-activated by YARS during cell emergence. An unbiased proteomic analysis confirmed the up-regulation of the Dream complex and of its targets during senescence escape.

## DISCUSSION

Senescence was initially described as a definitive proliferative arrest induced in response to telomere shortening, oncogenic expression or chemotherapy treatment. Whereas this pathway is most of the time definitive in primary cells, several studies have reported that this is not always the case in established cancer cells. Understanding how cells can enter a state of incomplete senescence and restart proliferation is an important question since this is certainly related to tumor dormancy and drug resistance, a major concern of chemotherapy treatment. We have recently described that the stability of chemotherapy-mediated senescence relies on specific tRNAs such as the tRNA-Leu-CAA and tRNA-Tyr-GTA (14). In addition, we also showed that their corresponding tRNA ligases, LARS and YARS are down-regulated during senescence and that their functions are necessary when cells restart proliferation. In this study, we extend these observations and describe non-conventional functions of YARS during senescence. As previously reported in other experimental conditions (20), we found that this tRNA ligase is present in the nucleus of colorectal and mammary cells. In association with Trim28, YARS binds the LIN9 promoter and support the loading of RNA Pol II. Down-regulating this ligase inhibits LIN9 expression, reduces proliferation and induces a senescent phenotype. During senescence escape, the re-activation of LIN9 expression by YARS promotes the emergence of proliferating cells.

Altogether, these results show that the ligase has non-conventional transcriptional functions during senescence and we speculate that this contributes to the stability of this proliferative arrest. It is important to note that the consequences of YARS inactivation depend on the cell line since its effects on senescence were observed in the MCF7 mammary cell line but not in LS174T colorectal cells. The ligase interacts with Trim28 and regulates cell proliferation in LS174T cells but its inactivation leads to senescence only in the mammary cell line. This heterogeneous response confirms our previous observation showing that different pools of tRNA ligases and tRNAs are expressed in senescent cells and that their effect depend on the experimental model. We speculate that the functions of YARS relies on three explanations that require further studies: the regulation of its nuclear translocation, the regulation of its transcriptional specificity and the specific functions of the Dream complex.

The main function of a tRNA ligase is to charge the tRNA with a cognate amino acid. Within YARS, a KKKLKK motif located between the amino acids 242 to 247 allows the interaction with the tRNATyr and the subsequent aminoacylation reaction. Importantly, this motif also serves as a nuclear localization signal but the contact with the import machinery relies on the accessibility of the KKKLKK sequence (26). As a consequence, the nuclear localization of YARS depends on the availability of the tRNATyr and on its binding to this sequence. When the expression of the tRNA decreases, the ligase interacts more efficiently with the import proteins and its presence in the nucleus increases. The expression of YARS and the tRNATyr, their interaction through the KKKLKK sequence, and the consequence on the nuclear localization of the ligase are probably different between MCF7 and LS174T cells, either during senescence induction or during cell emergence. Further studies are therefore necessary to clarify the nuclear translocation of YARS and understand its regulation in different cell lines and oncogenic context.

When present in the nucleus, the transcriptional function of YARS relies, at least in part, on its interaction with Trim28. This binding was observed in the two cell lines but the LIN9 promoter is certainly not the only target of these proteins. The RNA seq analysis performed in this study identified several cell cycle genes that could also be regulated by YARS and Trim28 and that could be involved in senescence. For instance, it will be interesting to determine if these two proteins regulate the Aurora-A promoter since we and others have already reported that this mitotic kinase plays an important role in chemotherapy resistance (27). Further studies are therefore required to identify the genes co-regulated by YARS and Trim28 that are involved in the regulation of senescence. In addition, it will be also necessary to clarify their transcriptional functions during this proliferative arrest. Our results indicate that both proteins are necessary for the loading of RNA pol

II on the LIN9 promoter and for the phosphorylation of its Ser2 site within the CTD domain. Interestingly, it has been previously reported that Trim28 regulates transcriptional elongation. In hypoxic breast cancer, Trim28 allows the recruitment of the cdk9 kinase on HIF-target genes, which then phosphorylates the CTD domain to allow gene elongation (28). However, it has also been shown that Trim28 induces RNA PolII pausing on the HSP70 promoter (29). This result was confirmed using ChIP-Seq analysis since 41% of the detected genes were regulated at the RNA polII pausing level. Further studies are therefore necessary to determine if YARS regulates the elongation and pausing functions of Trim28 on the LIN9 promoter.

The functions of Trim28 are complex since this protein also functions as a transcriptional repressor, interacting with the NuRD HDAC complex, the HP1 protein, and the SETDB1 histone methylase (30). It has been proposed that Trim28 modifies the local chromatin structure to form a stable heterochromatin state and induce transcriptional repression. This hypothesis is coherent with our observation that YARS down-regulation reduces the deposit of the H3K4me3 and H3K27ac activating marks on the LIN9 promoter. As stated above, we can speculate that these transcriptional and epigenetic functions of YARS depend on the cell line and promoters and that this might participate to the heterogeneity of the senescence response.

Further studies are also needed to characterize the role of YARS in regulating the Dream complex and how this might affect senescence stability. Recent results have already reported that the Dream complex is required for the induction of senescence. In response to the Ras oncogene, p130, E2F4, and DP1 bind to the MuvB-like proteins (LIN9, LIN37, LIN52, LIN54, and RBBP4) to repress cell cycle genes. Disrupting Dream association allows escape from Ras-mediated senescence (21) but it remains to be determined if this is also the case in response to chemotherapy. In addition, Kumari et al. have recently shown that the over-expression of LIN52, B-MYB, FOXM1, and to a lesser extent of LIN9 induce senescence bypass in human breast fibroblasts (31). By promoting LIN9 expression, we speculate that YARS may alter the composition of the Dream complex to regulate its transcriptional functions during senescence escape. This might lead to an abnormal expression of genes involved in the regulation of the G2/M phase of the cell cycle, which are known to be involved in senescence induction and escape (32). Further studies are therefore required to clarify the role of this pathway as the functions of LIN 9 and Dream during senescence remain largely unknown.

Altogether, these results describe non-canonical functions of YARS during senescence and senescence escape. During cell emergence, the ligase is present in the nucleus to activate LIN9 expression and probably the function of the Dream complex. Further experiments are needed to analyze these transcriptional functions and to determine if this could explain the heterogeneity of the senescence response.

## MATERIALS AND METHODS

### Cell Culture and senescence induction

MCF7 and LS174T cells were obtained from American Type Culture Collection (ATCC). Cells were cultured in Rosa Park Institut Medium 1640 (RPMI 1640 w/L Glutamine, Dutscher L0500-500) supplemented with 10% Fetal Bovine Serum (FBS, Eurobio CVFSVF00-01) without any antibiotics and at 37°C and 5% CO_2_. To induce senescence, MCF7 and LS174T cells were treated for 96 hours in RPMI medium supplemented with 3% of FBS. LS174T were treated with sn38 (5ng/ml, Tocris Bioscience 2684), and MCF7 were treated with doxorubicin (25ng/ml, Tocris Bioscience 2252). To promote senescence escape, cells were washed with phosphate buffered saline (PBS, Eurobio CS2PBS01K-BP) and stimulated with fresh RPMI medium supplemented with 10% FBS for 7 to 14 days as indicated.

### siRNA Transfection

Cells were transfected with 12.5nM or 50nM of a smart pool of 4 small interfering RNAs targeting LARS (ON-TARGETplus Human LARS (51520), L-010171-00-0005), YARS (ON-TARGETplus Human YARS (8565), L-011498-00-0005), CARS (ON-TARGETplus Human CARS (833), L-010335-01-0005), WARS (ON-TARGETplus Human WARS (7453), L-008322-00-0005), TRIM28 (ON-TARGETplus Human TRIM28 (10155), L-005046-00-0005), LIN9 (ON-TARGETplus Human LIN9 (286826), L-018918-01-0005) and prevalidated control siRNA (Dharmacon, D-001810-10-20) using DharmaFect-4 (Dharmacon, T-2004-03). Cells were transfected in medium containing 50% of OptiMEM (Fischer Scientific, 11524456) and 50% of RPMI 1640 medium supplemented with 10% FBS. After 24 hours, cells were washed with PBS and RPMI 1640 with 10% FBS was added.

### Cell Cycle Analysis

MCF7 or LS174T cells were washed with PBS and 250 000 to 500 000 cells were incubated with 300 uL of a solution A (30µg/ml of Trypsin, Sigma T0134, 3.4 mM Trisodique Citrate, Sigma C0909, 0.1% Igepal CA-630, Sigma I3021, 3mM Tetrahydrochloride spermine, Sigma S2876, 1mM Tris-aminométhane, Sigma T1378, pH = 7.6) for 10 minutes in the dark at room temperature. Then, 250 µL of solution B (0.5 mg/mL of Trypsin inhibitor, Sigma T9253, 0.1 mg/mL RNAse Type A, Sigma R4875, 3.4 mM Trisodique Citrate, Sigma C0909, 0.1% Igepal CA-630, Sigma I3021, 3mM Tetrahydrochloride spermine, Sigma S2876, 1mM Tris-aminométhane, Sigma T1378, pH = 7.6) was added for 10 minutes in the dark at room temperature. Finally, cells were incubated with 250 uL of solution C (0.6mM of propidium iodide, Sigma P4170, 3.3mM spermine tetrahydrochloride, Sigma S2876, 3.4 mM Trisodique Citrate, Sigma C0909, 0.1% Igepal CA-630, Sigma I3021, 3mM Tetrahydrochloride spermine, Sigma S2876, 1mM Tris-aminométhane, Sigma T1378, pH = 7.6) for 10 minutes à 4°C in the dark. Analysis was performed using a BD LSRII flow cytometer, recording 30 000 to 50 000 events per sample.

### SA-β-Galactosidase staining

Cells were fixed for 15 minutes at room temperature in 2% formaldehyde (Sigma, F8775). After fixation, cells were washed with PBS and incubated at 37°C (without CO2) with freshly-made senescence associated β-Gal (SA-β-Gal) staining solution (0.3 mg/ml of 5-bromo4-chloro-3indolyl β-D-galactoside, X-Gal Fermentas R0401, 40 mM citric acid, Sigma C-0759, 40 mM sodium phosphate, Sigma S5011, (stock solution, 400 mM citric acid, 400 mM sodium phosphate, pH = 6), 5 mM potassium ferricyanide, Sigma 244023, 5 mM ferrocyanide, Sigma P3289, 150 mM NaCl, Sigma S5886 and 1.5 mM MgCl2, Sigma M8266) for 16 to 20 hours. Positive SA-β-Galactosidase cells display a perinuclear precipitation of blue dye, which can be observed with standard light microscopy.

### RT-qPCR

Cells were washed with PBS and mRNAs were extracted using NucleoZOL (MACHEREY-NAGEL, 740404.200) according to the manufacturer’s instructions but the step with water was repeated twice to avoid contamination by genomic DNA. Quantification of mRNA was performed by nanodrop. Reverse transcription was performed after incubation of mRNA with Random Primers (ThermoFisher Scientific, 48190011) for 5 minutes at 70°C, then a mix containing M-MLV buffer, dNTPs, and M-MLV reverse transcriptase (ThermoFisher Scientific, 28025013) was added, and samples were incubated for 1 hour at 37°C.

Quantitative PCR was performed using the Maxima SYBR Green/ROX qPCR master mix (ThermoFisher Scientific, K0223) with a mix of forward and reverse primer (with a final concentration of 0.5µM). Samples were amplified 40 cycles using Quanti Studio 3. Analysis was performed using the comparative CT method (2^(ΔCt)), according to the expression of three endogenous housekeeping gene TBP, PPIA and EEF1A1 or βActin. All primers sequences are provided below.

### Flow cytometry

Cells were recovered using PBS-EDTA 2mM, then washed with PBS-BSA 2% twice, and 250 000 cells were incubated for 15 minutes in the dark at room temperature with 200 ng of APC anti-CD47 (eBiosciences, 17-0479-42) or 200 ng of APC mouse IgG1K isotype control (eBiosciences, 17-4714-42). Then cells were washed with PBS-BSA 2% and analysis was performed using a BD LSRII flow cytometer, recording 30 000 to 50 000 events per sample.

### Western Blot

Cells were lysed with FASP Buffer containing inhibitors of protease and phosphatase (0.1 M Tris-HCL, 4% SDS, pH=7.6 and 10 µg/ml aprotinin, 10 µg/ml leupeptin, 10 µg/ml pepstatin, 1 mM Na3VO4, 50 mM NaF) and then sonicated and boiled for 10 min. Protein concentrations were determined using the BCA protein assay kit (Pierce, 23227). Equal quantity of proteins were separated on a SDS polyacrylamide gel and transferred to a PVDF membrane. Following a 1 hr incubation in 5% milk, Tris-buffered saline (TBS), and 0.1% Tween 20, membranes were incubated overnight at 4 °C with the following primary antibodies: p-eIF2a S51 (1/1000, Cell Signaling, 9721), YARS (1/1000, SantaCruz, sc166741), Lamin A/C (1/1000, SantaCruz, sc7293), α-Tubulin (1/1000, SantaCruz, sc5286), Actin (1/1000, SantaCruz, sc8432), TRIM28 (1/1000, Abcam, ab10484), CARS (1/1000, SantaCruz, sc390230), LARS (1/1000, Cell signaling, 35509), WARS (1/1000, Abcam, ab109213), p21waf1/cip1 (1/1000, Cell Signaling 2947), Hsc70 (1/1000, SantaCruz, sc7298). Then membranes were washed three times with TBS with 0.1% Tween 20 for 5 minutes at room temperature and incubated with the secondary antibodies listed below: Anti-rabbit IgG, HRP-linked antibody (1/3000, Cell Signaling, 7074), Anti-mouse IgG, HRP-linked Antibody (1/3000, Cell Signaling, 7076) for 45 minutes at room temperature. Visualization was performed by chemiluminescence with a Fusion Solo apparatus (Vilber).

### Immunofluorescence

Cells were fixed for 15 minutes at room temperature in 2% formaldehyde (Sigma, F8775) and washed 3 times with PBS. Then, 1ml of Methanol 70% was added and samples were conserved at 4°c for at least 12 hours. Cells were then washed 3 times with PBS 0.02% Tween and incubated with PBS 2% BSA, 10 minutes at room temperature. Then cells were incubated overnight à 4°C with the following antibodies : YARS (SantaCruz, sc166741), ki67 (Cell Signaling, 9449), p21 (Cell Signaling, 2947), Rabbit IgG Isotype Control (Cell Signaling, 3900), Mouse IgG Isotype Control (Cell Signaling, 5415). Cells were then washed 3 times with PBS 0.02% Tween and incubated with PBS 2% BSA, 10 minutes at room temperature. Then, cells were incubated with the following secondary antibodies diluted 1/200 : Goat anti-Mouse IgG secondary antibody (Alexa 488, Invitrogen-Molecular Probes : A-A11001), Goat anti-Mouse IgG secondary antibody (Alexa 568, Invitrogen-Molecular Probes : A-11004) Goat anti-Rabbit IgG secondary antibody (Alexa 488, Invitrogen-Molecular Probes : A-11008), Goat anti-Rabbit IgG secondary antibody (Alexa 568, Invitrogen-Molecular Probes : A-11011) for 1 hour at room temperature in the dark. Cells were then washed 3 times in PBS 0.02%Tween, and covered with Anti-Fading reagent with DAPI (Invitrogen Prolong R. Gold anti-fade reagent with DAPI P36935). The slides were subsequently analyzed using confocal microscopy on the Sciam microscopic platform (https://sfricat.univ-angers.fr/fr/plateformes/sciam.html). Quantification analysis was performed using the ImageJ software.

### 53BP1 Staining

Cells were washed with PBS, fixed for 20 minutes at room temperature in 10% formalin and washed 3 times with PBS-Glycine (50mM). Then, 0.5ml of permeabilization buffer (NaCl 10mM, Sucrose 300mM, Pipes 10mM, MgCl2 5mM, 5% Triton 100X) was added and samples were incubated 10min and washed twice with permeabilization buffer without triton. After this, cells were incubated with IF buffer (NaCl 130mM, 13.2 mM Na2HPO4, 3.5mM NaH2PO4, 0.1% bovine serum albumin, 0.2% Triton X-100 and 0.05% Tween 20) containing 10% of goat serum for 1 hour at room temperature. Then cells were washed three times with IF buffer and incubated overnight at 4°C with the following antibodies : 53BP1 (abcam, ab36823) or Rabbit IgG Isotype Control (CellSignaling, 3900). After this, cells were washed 3 times with IF buffer and incubated with the following secondary antibody: Goat anti-Rabbit IgG secondary antibody (Alexa 568, Invitrogen-Molecular Probes : A-11011) for 1 hour at room temperature in the dark. Cells were then washed 3 times with IF buffer, and covered with Anti-Fading reagent with DAPI (Invitrogen Prolong R. Gold antifade reagent with DAPI P36935). Slides were then analyzed by microscopy. Quantification analysis was performed using the ImageJ software.

### Co-immunoprecipation

Cells were lysed in Lysis Buffer pH 7.4 (20mM Tris-HCl, 150mM NaCl, 1mM EDTA, 1% Triton X100, 1 mM PMSF, 10 µg/ml aprotinin, 10 µg/ml leupeptin, 10 µg/ ml pepstatin, 1 mM Na3VO4, 50 mM NaF). Lysats were pre-cleared with 20µL of magnetics beads (FischerScientific, 13484209) for 2 hours at 4°C. Then lysats were incubated with 1µg of the following antibodies : YARS (SantaCruz, sc166741), TRIM28 (Abcam, ab10484), Rabbit polyclonal IgG (Abcam, ab171870), Mouse monoclonal IgG (Abcam, ab18413), overnight at 4°C. 20µL of magnetics beads were then added for 2 hours at 4°C. After two washes with lysis buffer, western blot was performed as described above, except that the revelation was performed using Anti-goat IGG Veriblot for IP secondary antibody (Abcam, ab157532).

### Proximity Ligation Assay

Cells were fixed for 15 minutes at room temperature in 2% formaldehyde (Sigma, F8775) and washed 3 times with PBS. Then, 1ml of Methanol 70% was added and samples were conserved at 4°C for at least 12 hours. Samples were then washed 3 times with TBS 0.05% Tween 20 and the detection was performed using the Naveniflex kit (NF.100.MR) according to the manufacturer’s instructions. The following primary antibodies were used : YARS (SantaCruz, sc166741), TRIM28 (Abcam, ab10484), Rabbit polyclonal IgG (Abcam, ab171870), Mouse monoclonal IgG (Abcam, ab18413). At the step 7, the Anti-Fading reagent with DAPI (Invitrogen Prolong R. Gold antifade reagent with DAPI P36935) was used. Slides were then analyzed by microscopy or confocal microscopy.

### Nuclear and cytoplasmic protein extraction

Cells were lysed with Harvest buffer (Hepes 10 mM, NaCl 50 mM, EDTA 10 mM, Sucrose 0.5M, Triton X100 0.5%), incubated for 5 min on ice and then cell lysates were centrifuged for 10 min at 800g. The supernatant was removed and centrifuged at 10000 g for 15 min at 4°C to get the cytosolic fraction. The nuclear pellet was washed twice with buffer A (Hepes 10 mM, EGTA 0.1 mM, EDTA 0.1 mM, KCl 10 mM) and centrifuged at 800g for 10 min at 4°C. The final pellet was resuspended with High Salt buffer (Hepes 25 mM, NaCl 0.4M, EDTA 0.2 mM, MgCl2 1.5 mM, NP40 1%) and incubated for 60 min on rotation at 4°C. Finally, nuclear extracts were centrifuged at 10,000g for 15 min at 4°C. The supernatants, containing nuclear proteins, were collected and each fraction was used for Western blotting.

### Mass Spectrometry and ATACseq/RNAseq analysis

As indicated in the corresponding figures, growing or senescent LS174T and MCF7 cells were used. Cells were treated with chemotherapy for 96 hours in RPMI medium supplemented with 3% FBS to induce senescence. Control cells were grown in the same conditions but in the absence of treatment. After 96hr, cells were recovered, centrifuged and 100 00 cells were used for ATACseq experiments, between 600 000 and 1,5 millions cells for RNA sequencing. ATACseq results correspond to two independent experiments, RNA sequencing results to three. Samples were sent to Active motif for library preparation and sequencing analysis. For RNA sequencing, total RNA was isolated from the cells using the Qiagen RNeasy Mini Kit (Qiagen, cat# 74104). For each sample, 2 ug of total RNA was then used in Illumina’s TruSeq Stranded mRNA Library kit (Cat# 20020594). Libraries were sequenced on Illumina NovaSeq 6000 as paired-end 42-nt reads. Sequence reads were analyzed with the STAR alignment – DESeq2 software pipeline. For for ATACseq experiments, cryopreserved cells were thawed in a 37°C water bath, pelleted, washed with cold PBS, and tagmented as previously described (Buenrostro et al. Nature Methods 2013), with some modifications based on (Corces et al. Nature Methods 2017). Briefly, cell pellets were resuspended in lysis buffer, pelleted, and tagmented using the enzyme and buffer provided in the Nextera Library Prep Kit (Illumina). Tagmented DNA was then purified using the MinElute PCR purification kit (Qiagen), amplified with 10 cycles of PCR, and purified using Agencourt AMPure SPRI beads (Beckman Coulter). Resulting material was quantified using the KAPA Library Quantification Kit for Illumina platforms (KAPA Biosystems), and sequenced with PE42 sequencing on the NovaSeq 6000 sequencer (Illumina). Reads were aligned using the BWA algorithm (mem mode; default settings). Duplicate reads were removed, only reads mapping as matched pairs and only uniquely mapped reads (mapping quality >= 1) were used for further analysis. Alignments were extended in silico at their 3’-ends to a length of 200 bp and assigned to 32-nt bins along the genome. The resulting histograms (genomic “signal maps”) were stored in bigWig files. Peaks were identified using the MACS 2.1.0 algorithm at a cutoff of p-value 1e-7, without control file, and with the –nomodel option. Peaks that were on the ENCODE blacklist of known false ChIP-Seq peaks were removed. Signal maps and peak locations were used as input data to Active Motifs proprietary analysis program, which creates Excel tables containing detailed information on sample comparison, peak metrics, peak locations and gene annotations. For differential analysis, reads were counted in all merged peak regions (using Subread), and the replicates for each condition were compared using DESeq2.

For proteomic analysis, growing or senescent LS174T and MCF7 cells were used as described above and analyzed by the proteomic platform of the ICO cancer center. Briefly, cell extracts were recovered and each sample was analyzed as previously described using a a data-independant acquisition (DIA)-MS method (see the (12,14) references for a complete description of the protocol). Data extraction of the DIA runs was performed by PeakView, MarkerView (version 1.2, Sciex) and Spectronaut (Biognosys, v17) were used for signal normalization, and differential abundance was tested by applying a t-test at the protein level. All data were analyzed using the ToppGene software (toppgene.cchmc.org/) to perform functional enrichment according to Chen et al., Nucleic Acids Res. 37, 305-311.

### Chromatin Immuno-precipitation (ChIP)

Cells were cross-linked with 1% formaldehyde (Sigma Aldrich) for 10 minutes at room temperature. Cross-linking was stopped by adding 0.125 mol/L glycine for 5 minutes. Cells were washed three times with cold PBS. Then cells were scraped and washed three times with cold PBS. Pellets were resuspended in 1 mL of lysis buffer (5 mM PIPES, 85mM KCl, and 0.5% NP40). Then samples were mixed using vortex every 2 minutes for 30 seconds, during 15 minutes at 4°C. After this, cells were centrifuged for 10 minutes, 16 000g at 4°C and pellets were resuspended with 500 µL of sonicating buffer (10 mM EDTA, 1% SDS, and 50 mM Tris-EDTA, pH 8). Samples were sonicated (11 cycles of 30s of sonication and 30s on ice) to obtain reverse cross-linked DNA fragments with a size of 500-200 bases. Supernatants were diluted 10 times with IP buffer (0.01% SDS, 1.1%Triton X-100, 1.2 mM EDTA, 16.7 mM Tris-HCl (pH =8.0), and 167 mM NaCl). For each condition, between 8 and 12 µg of chromatin was used (chromatin quantity was normalized between conditions and concentration was estimated based on the reverse cross-linked chromatin). The chromatin was pre-cleared with 25 µl of beads for 2 hours on rotation at 4°C. Magnetics beads (FischerScientific, 13484209) were coated during the same time with 3 µg of the following antibodies: TRIM28 (Abcam, ab10484), YARS (SantaCruz, sc166741), RNA polymerase II (SantaCruz, sc47701), RNA polymerase II CTD repeat YSPTSPS phosphor Ser2 (Abcam, ab5095), H3K27 acetylated (Abcam, ab4729), H3K4me3 (Abcam, ab8580), Rabbit polyclonal IgG (Abcam, ab171870), Mouse monoclonal IgG (Abcam, ab18413). Each pre-cleared sample was then incubated with 45 µl of coated magnetic beads and DTT (1 mM) and BSA (10 µg/ml) added at the last moment. After overnight incubation at 4°C, the beads were washed successively with 1.5 ml of TSE1 Buffer (1%Triton X-100, 150 mM NaCl, 20 mM Tris-HCl, pH 8.1, 0.1% SDS, and 2 mM EDTA), 1.5mL of TSE2 buffer (1%Triton X-100, 500 mM NaCl, 20 mM Tris-HCl, pH 8.1,0.1% SDS, and 2 mM EDTA), and 1.5mL of TSE3 buffer 1% NP40, 1% sodium deoxycholate, 250mM LiCl, and 10mM Tris-HCl, pH 8.1). Following two washes in TE buffer (10 mM Tris-HCl and 1 mM EDTA), samples were eluted with 300 µL of fresh elution buffer (1% SDS and 0.1M NaHCO3). The cross-link was reversed by adding 24µL of NaCl (5 M) and 6µL of EDTA (0.5 M) followed by overnight incubation at 65 °C. DNA was purified using a High Pure PCR Template Preparation Kit (Roche, 11796828001) and q-PCR was performed as describe above. Data was normalized using the ‘‘Percent Input Method,’’ which means that data are normalized for background according to the Input sample (10% starting chromatin), the amount of chromatin used per ChIP. Values represent the percentage (%) of DNA immunoprecipitated according to the Input signal. The primers used for qPCR analysis are listed as a supplementary file.

### Statistical analysis

Differences were analyzed using a non-parametric test (Mann-Whitney or Kolmogorov-Smirnov for normalized data). * p<0.05, ** p<0.01 and *** p<0.001. When no stars are indicated on the graph it means that differences were not significant.

## Abbreviations

CIS: Chemotherapy-Induced Senescence
AARS: aminoacyl-tRNA synthetase
YARS: cytoplasmic Tyrosyl (Y) tRNA synthetase
LARS: cytoplasmic Leucyl (L) tRNA synthetase

## Acknowledgements

This work was supported by grants from the Ligue Contre le Cancer (Comité du Maine et Loire, du Finistère, de la Loire Atlantique) and the Rotary Club (Maine et Loire).

---

**Supplementary Figure 1: CARS, LARS and WARS are not involved in senescence induction in MCF7 and LS174T cells**

A. Growing cells were transfected with the indicated siRNAS, cell extracts were recovered after fours days and analyzed by western blot using the indicate antibodies (n=4 except WARS n=2).

B. Representative images of SA-β galactosidase staining 96 hours after the transfection of MCF7 ans LS174T growing cells with the indicated siRNAs (n=2 for each cell line except YARS n=4).

C. Growing LS174T cells were transfected with the indicated siRNAs as described above. After four days, the expression of the p15, p21 and p16 mRNAs was evaluated by RT-qPCR analysis (n=4).

D. MCF7 cells were treated with doxorubicin for 96 hours and FACS analysis was performed to analyze CD47 expression (n=4 * p<0.05).

E. MCF7 cells were treated with doxorubicin for 96 hours, then cells were fixed and stained with antibody directed against 53BP1. Results were analyzed by fluorescence microscopy and foci counting (n=2).

**Supplementary Figure 2: Senescence induction in LS174T and MCF7 cells following chemotherapy treatment**

A. Growing MCF7 and LS174T cells were treated respectively with doxorubicin or sn38 for 96 hours then FACS analysis was performed to analyze the cell cycle profile of the indicated cells (MCF7 n=3, LS174T n=4 * p<0.05).

B. Representative images of SA-β galactosidase staining 96 hours after doxorubicin or sn38 treatment of MCF7 or LS174T growing cells (n=3 for each cell line).

C. Growing MCF7 and LS174T cells were treated respectively with doxorubicin or sn38 for 96 hours, then the expression of the p15 and p21 mRNAs was evaluated by RT-qPCR analysis (n=4 for each cell line, * p<0.05).

D. Growing MCF7 and LS174T cells were treated respectively with doxorubicin or sn38 for 96 hours, then the expression of p21 was evaluated by western blot analysis (n=2 for each cell line).

E. Growing MCF7 and LS174T cells were treated with doxorubicin or sn38 for 96 hours, then the expression of the indicated mRNAs was evaluated by RT-qPCR analysis (n=4 for each cell line * p<0.05).

F. Growing MCF7 and LS174T cells were transfected with a control siRNA or a siRNA directed against YARS. Cells were fixed and stained with antibody raised against YARS. Results were analyzed by fluorescence microscopy.

**Supplementary Figure 3: Specificity of the Proximity-Ligation-Assay**

A. Total cell extracts were recovered from growing MCF7 and LS174T cells and the association of endogenous TRIM28 and YARS was analyzed by co-immunoprecipitation (n=2 for each cell line).

B. Growing MCF7 and LS174T cells were transfected with a control siRNA or a siRNA directed against YARS. Then, TRIM28 and YARS interaction was analyzed by Proximity-Ligation-Assay and analyzed by fluorescence microscopy (MCF7 n=2, LS174T n=1).

C. Growing MCF7 and LS174T cells were transfected with a control siRNA or a siRNA directed against YARS. Proximity-Ligation-Assay was performed using either YARS antibody alone (bottom) or YARS and TRIM28 antibodies combined with rabbit or mouse Ig control antibodies (n=1 for each cell line).

**Supplementary Figure 4 : TRIM28, YARS and LIN9 siRNAs and ChIP specificity**

A. Senescent MCF7 and LS174T cells were transfected with a control siRNA or a siRNA directed against TRIM28 for 24h. After three days, samples were recovered and TRIM28 expression was evaluated by western blot (n=3 for each cell line).

B. Growing MCF7 cells were transfected with a control siRNA or a siRNA directed against YARS or TRIM28 for 24h. Chromatin was recovered 3 days after depletion, then the binding of the indicated proteins was analyzed by ChIP experiments (n=4, * p<0.05).

C. Growing MCF7 and LS174T cells were transfected with a control siRNA or a siRNA directed against LIN9 for 24h. After three days, the expression of LIN9 mRNA was evaluated by RT-qPCR analysis (n=4 for each cell line, * p<0.05).

D. Growing MCF7 and LS174T cells were transfected with a control siRNA or a siRNA directed against LIN9 for 24h. After three days, the expression of the indicated mRNAs was evaluated by RT-qPCR analysis (n=4 for each cell line, * p<0.05).

