## Supplementary figures and images for "The YARS Tyrosyl tRNA Synthetase Regulates Senescence Induction and Escape Through the Transcriptional Control of LIN9, a Member of the DREAM Complex"

### Supplementary Figures

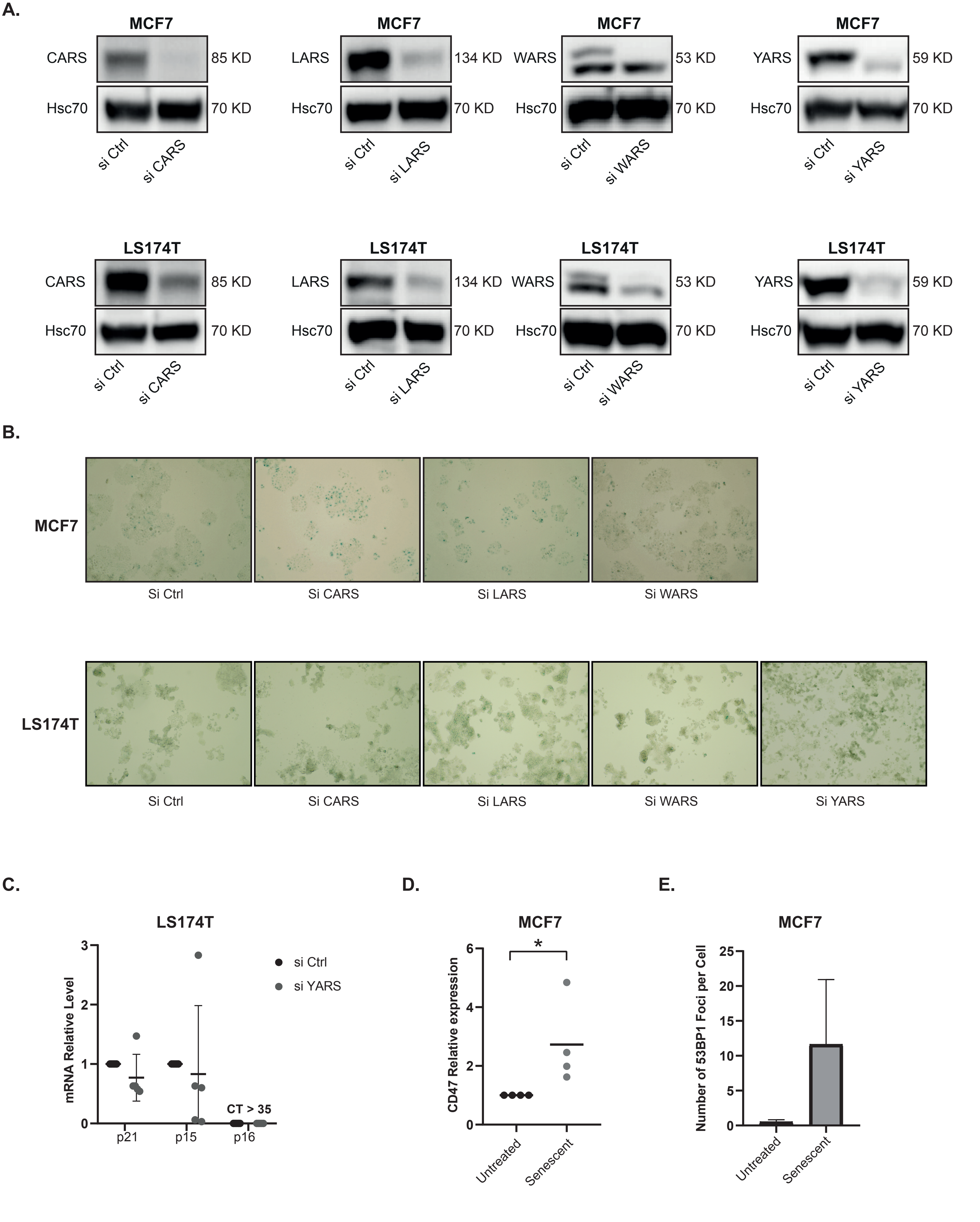


Supplementary Figure 1

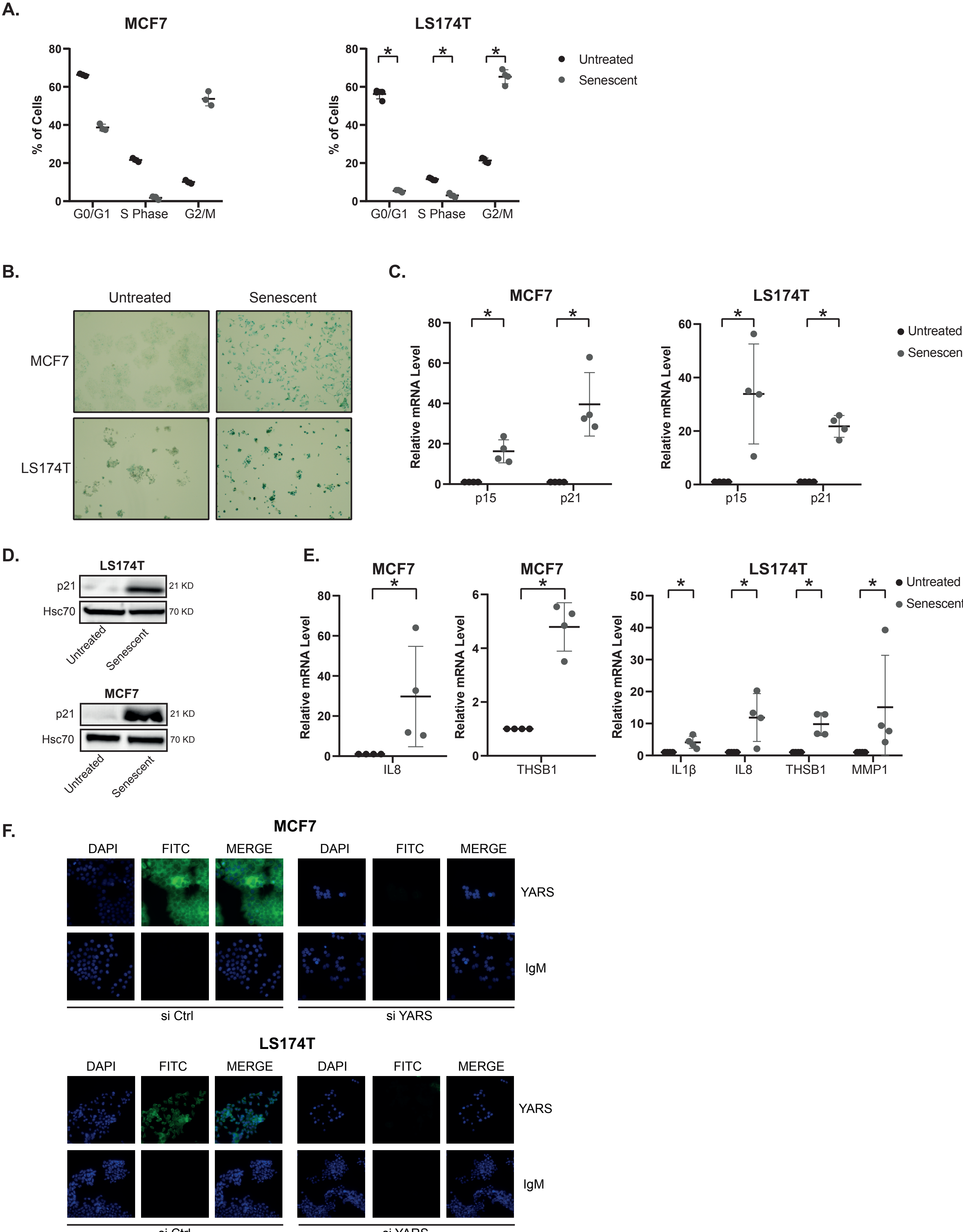


Supplementary Figure 2

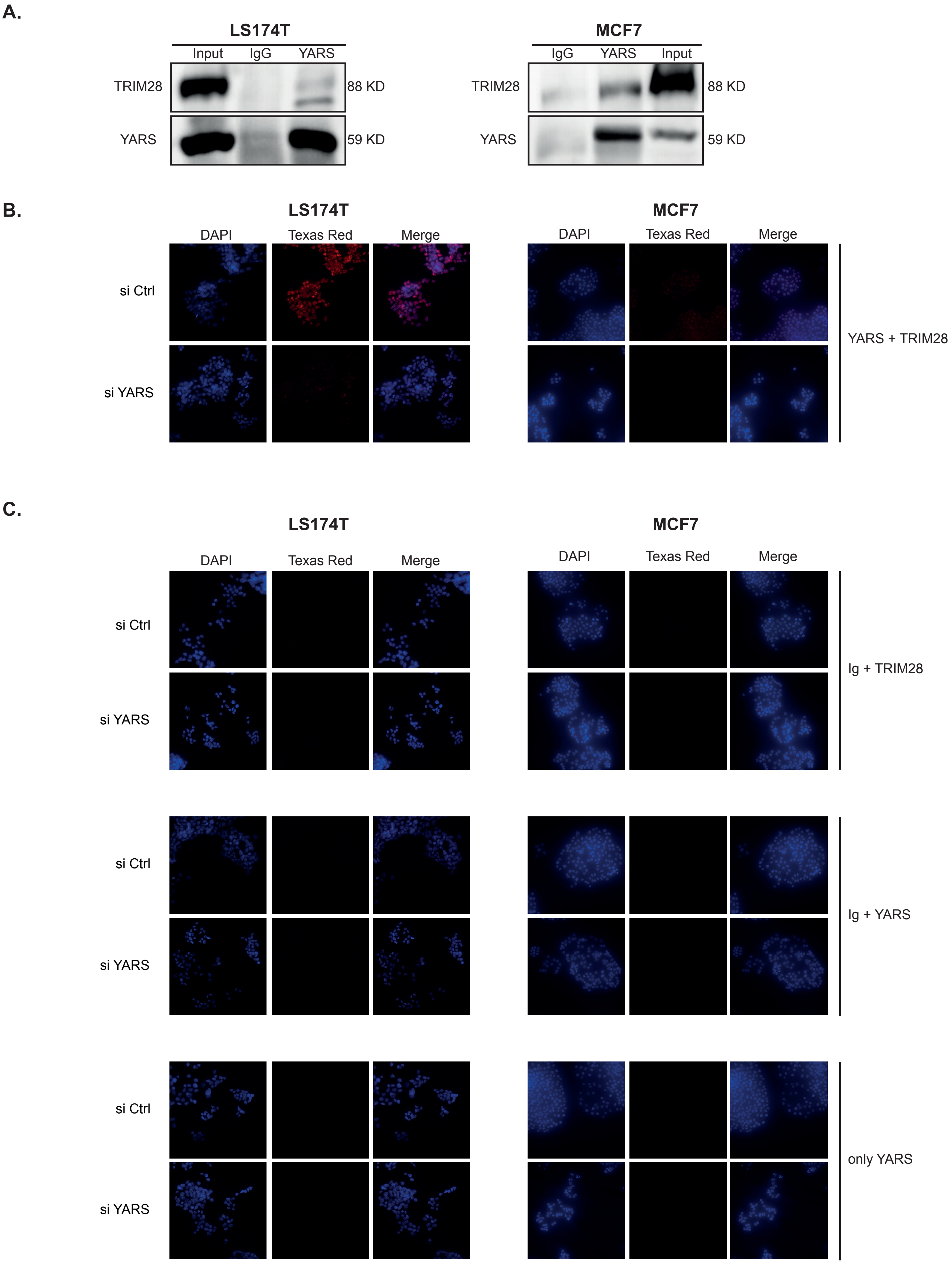


Supplementary Figure 3

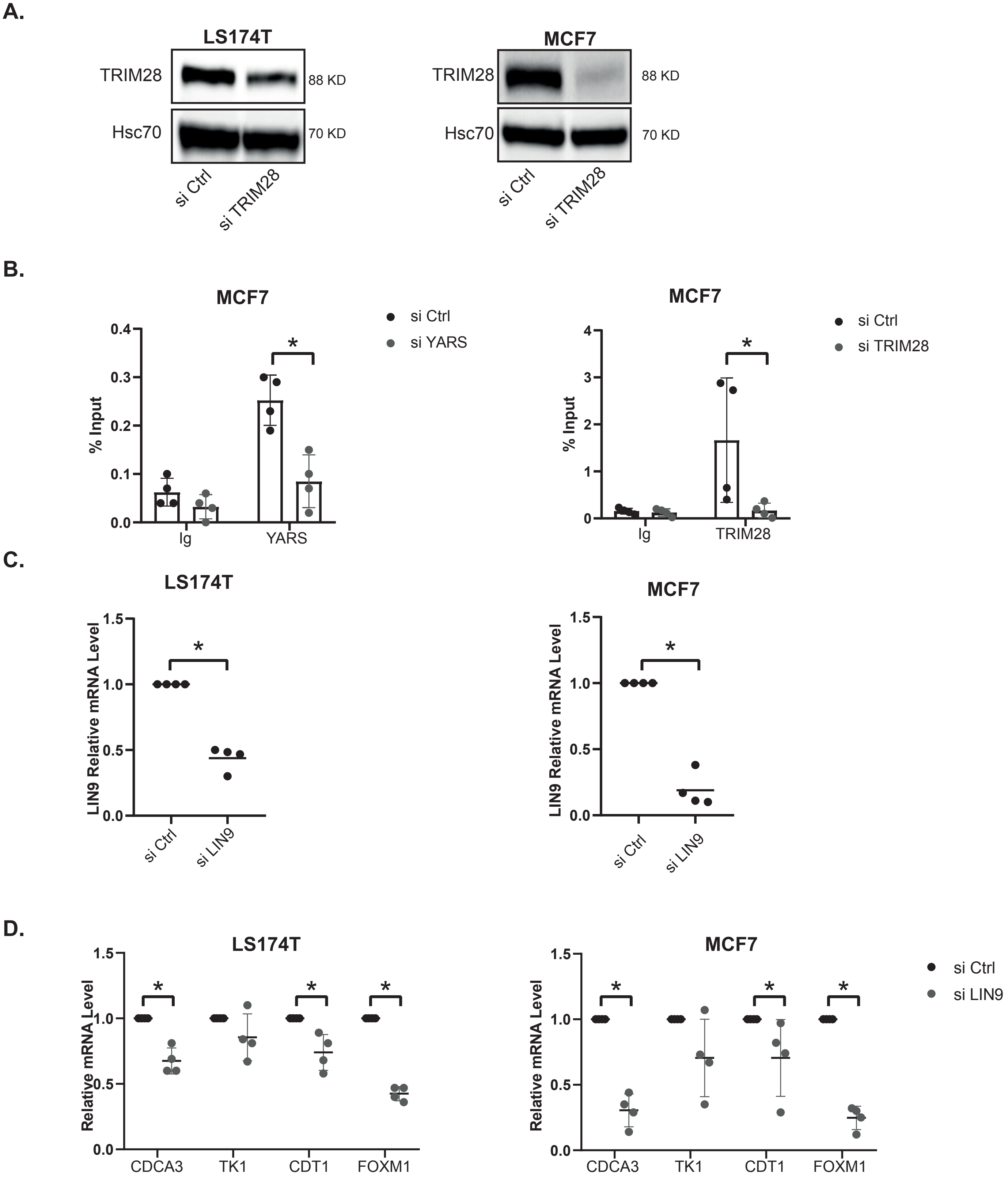


Supplementary Figure 4
